## Supplementary figures and images for "The actin module of endocytic internalization in *Aspergillus nidulans*: a critical role of the WISH/DIP/SPIN90 family protein Dip1"

### Supplemental figures

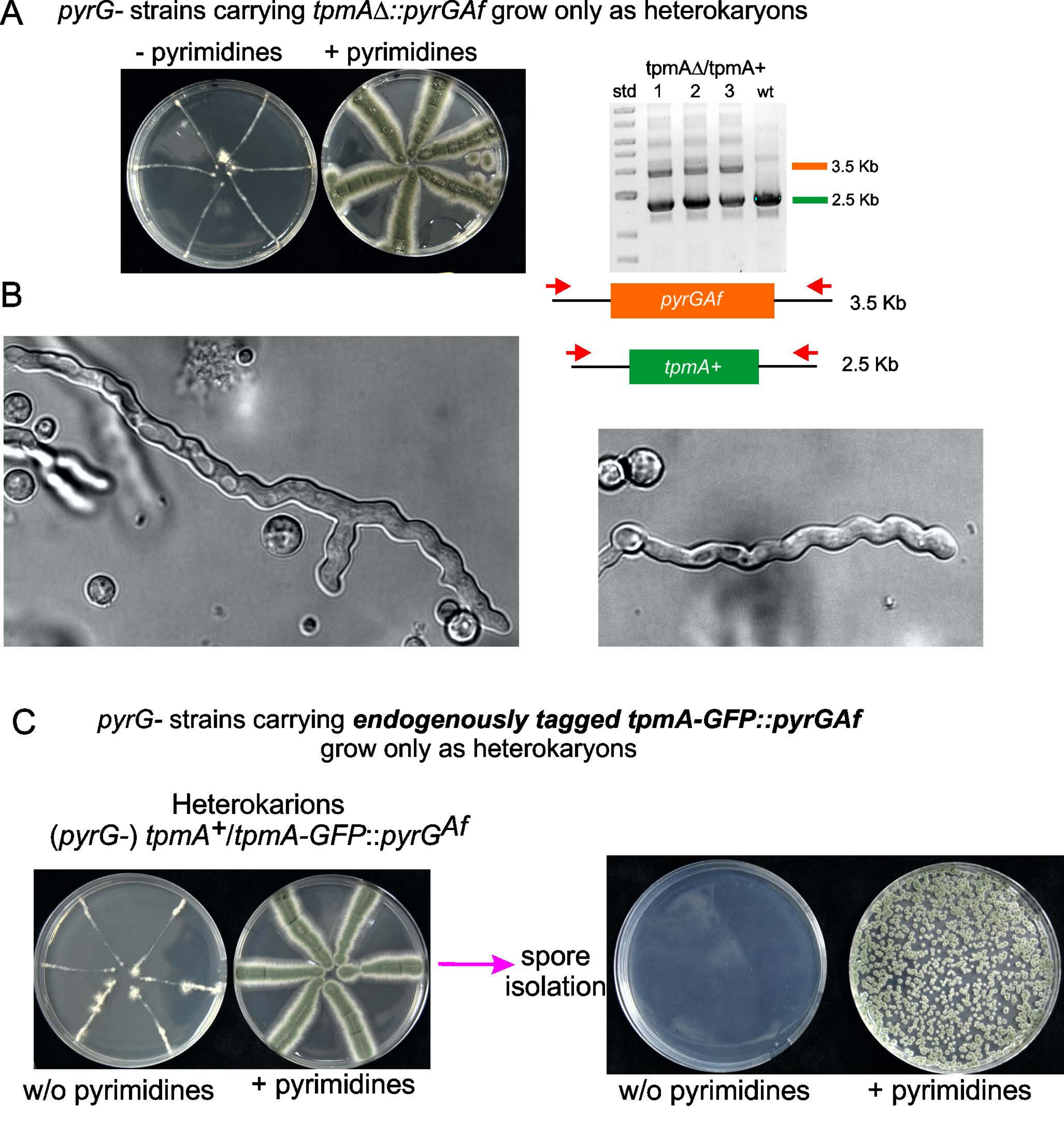

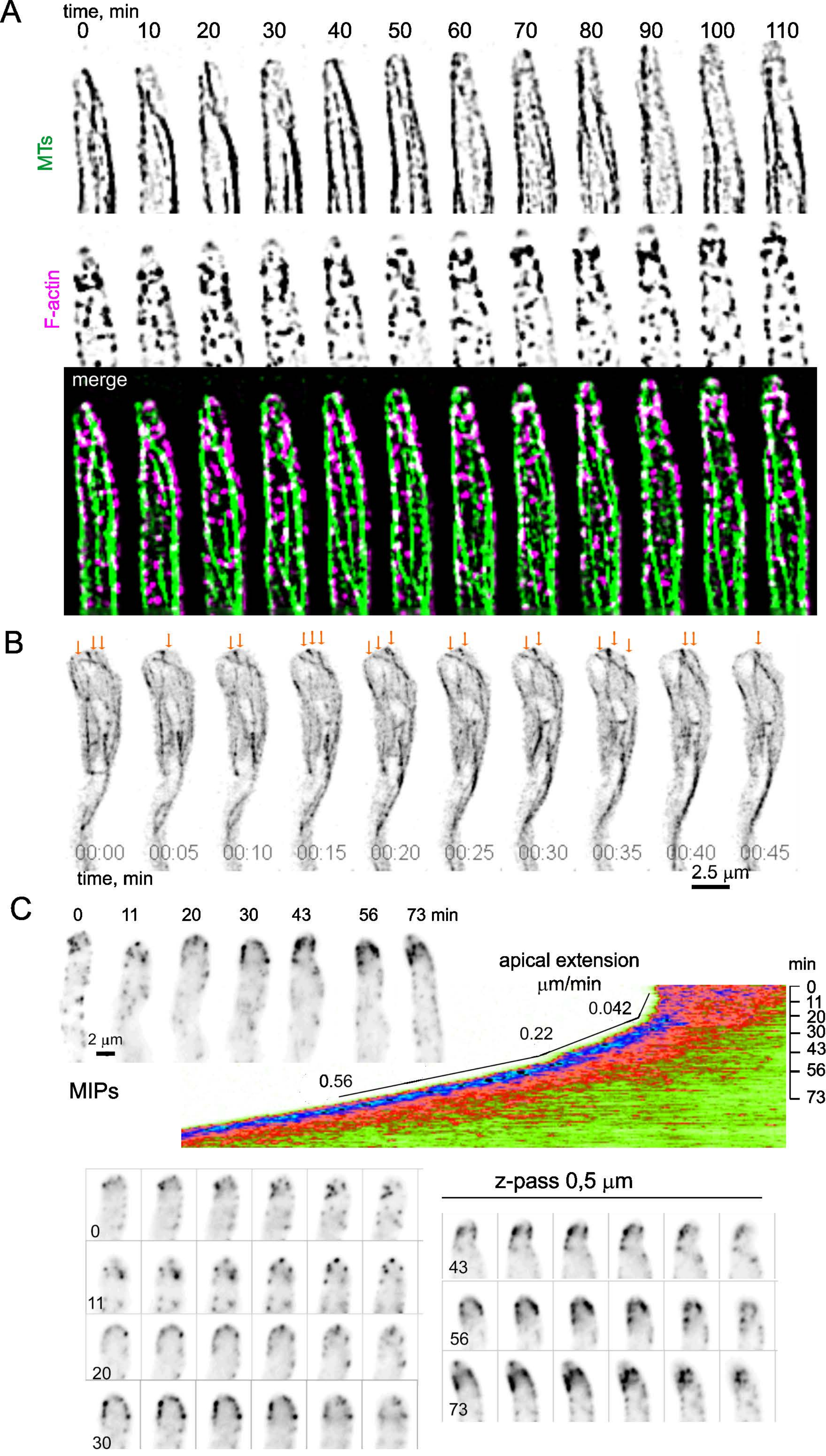

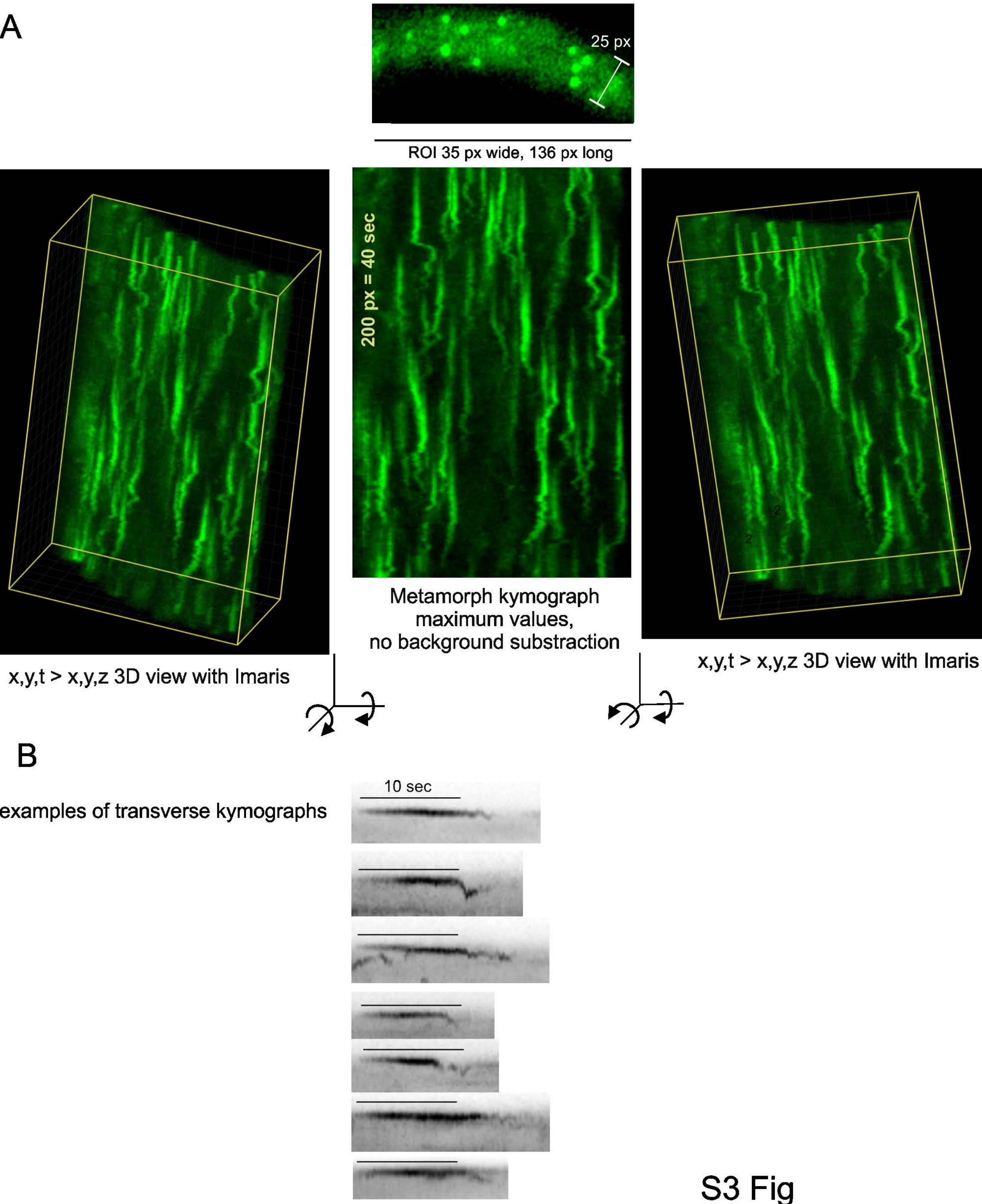

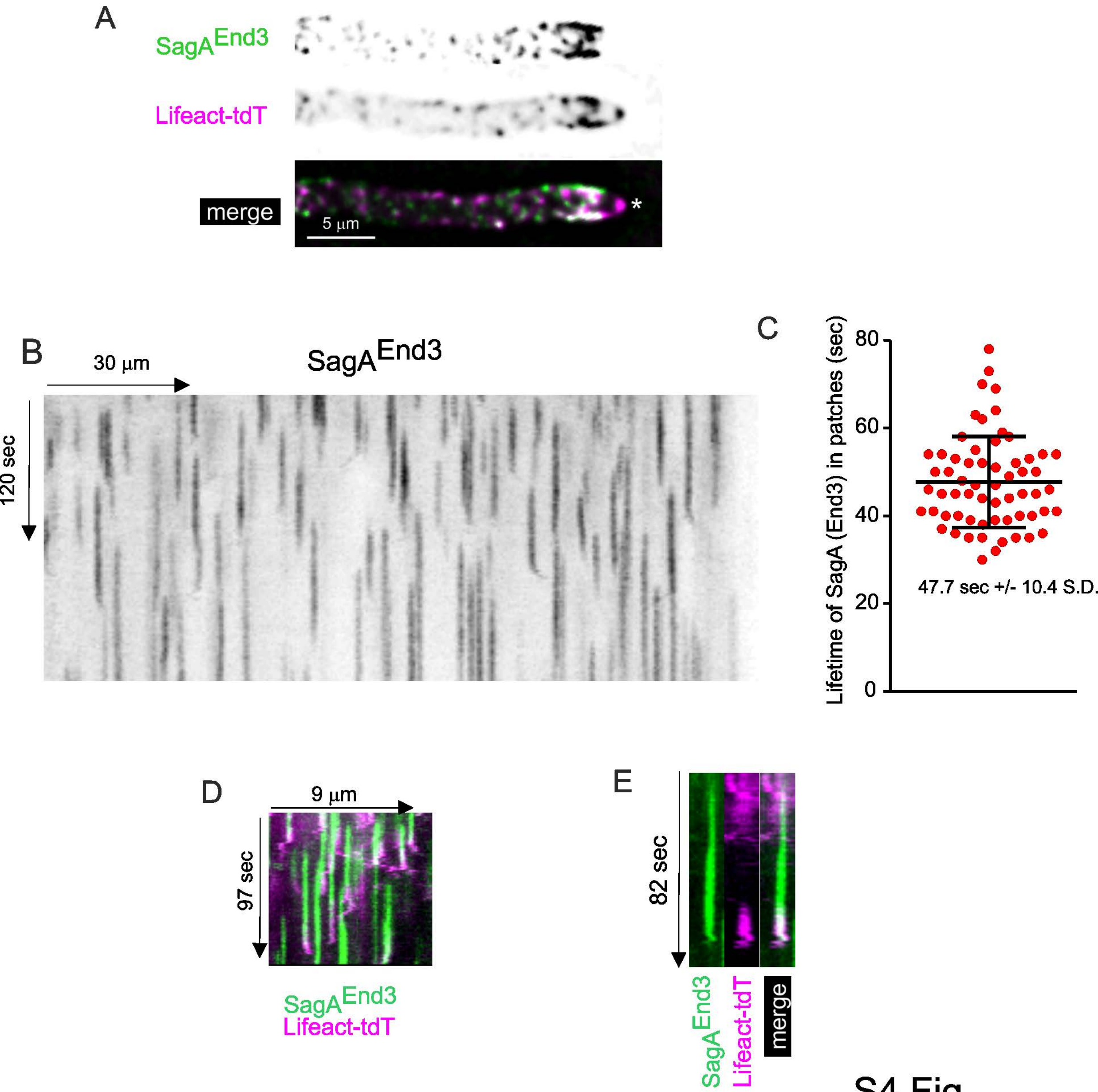

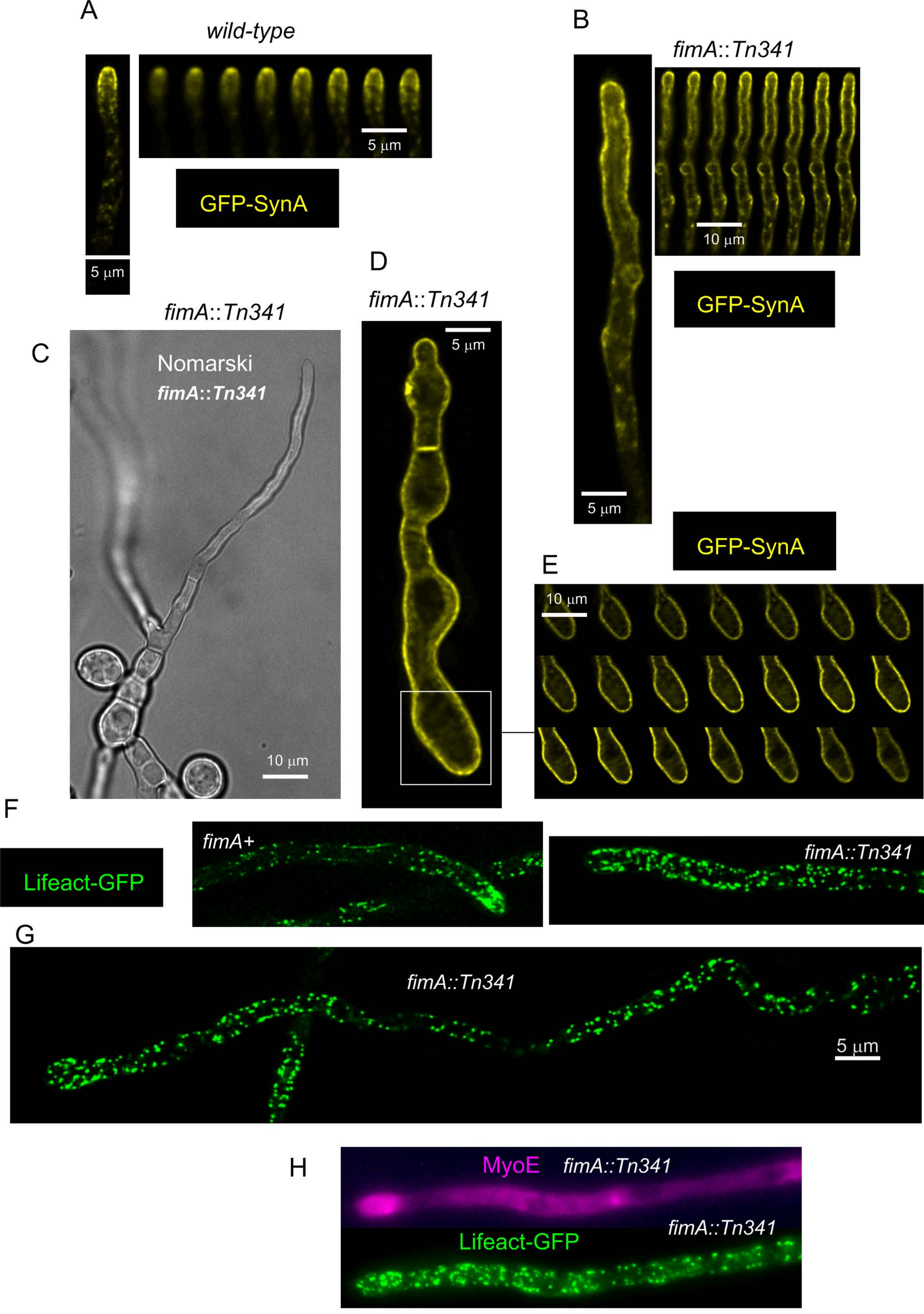

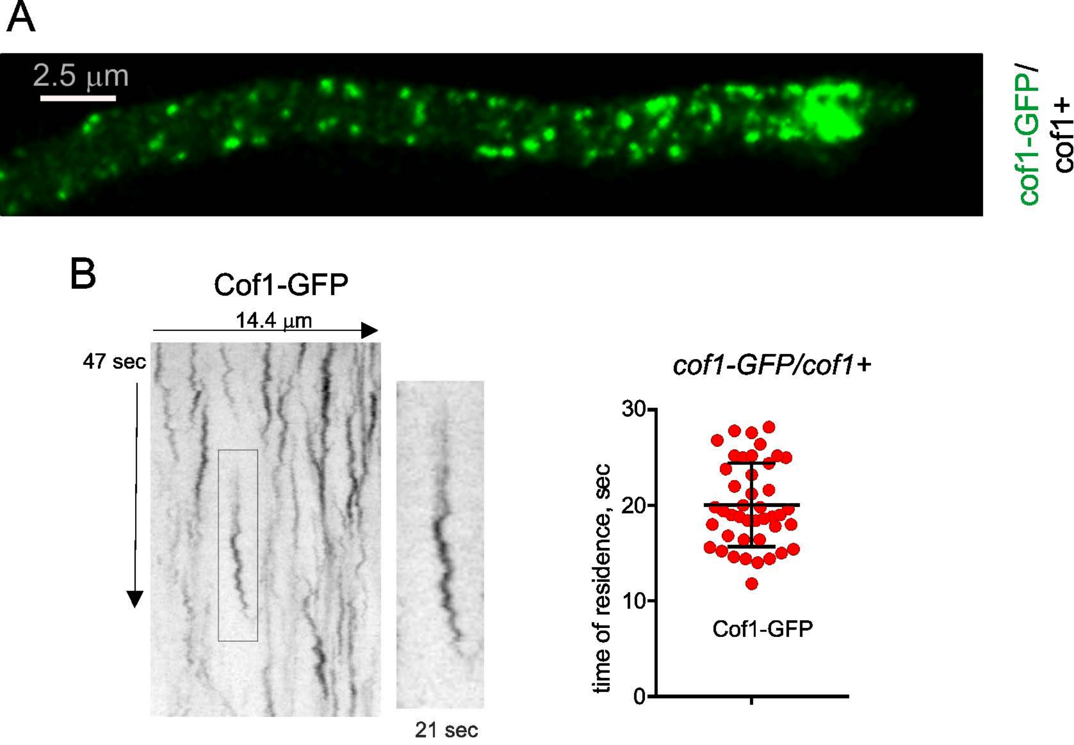

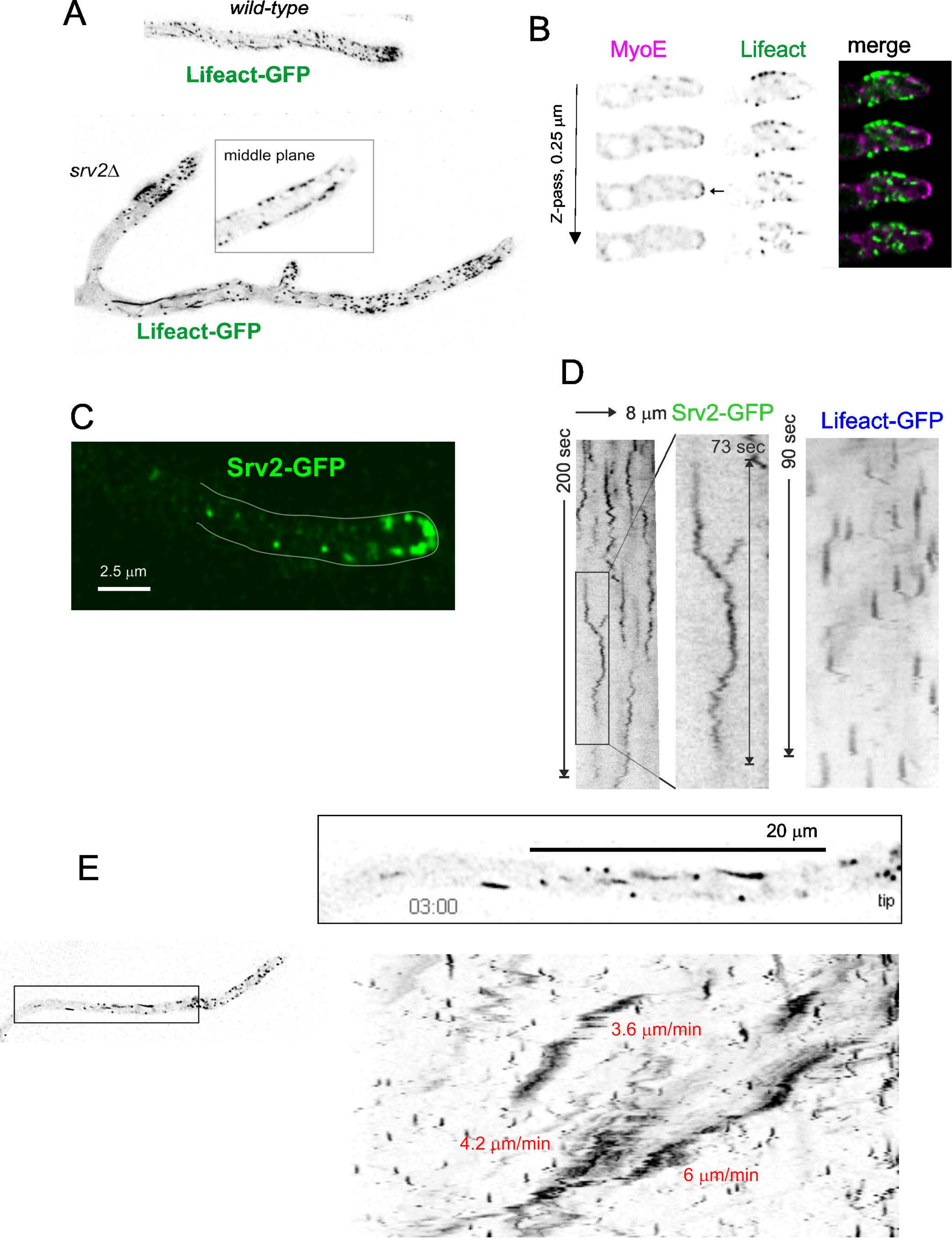

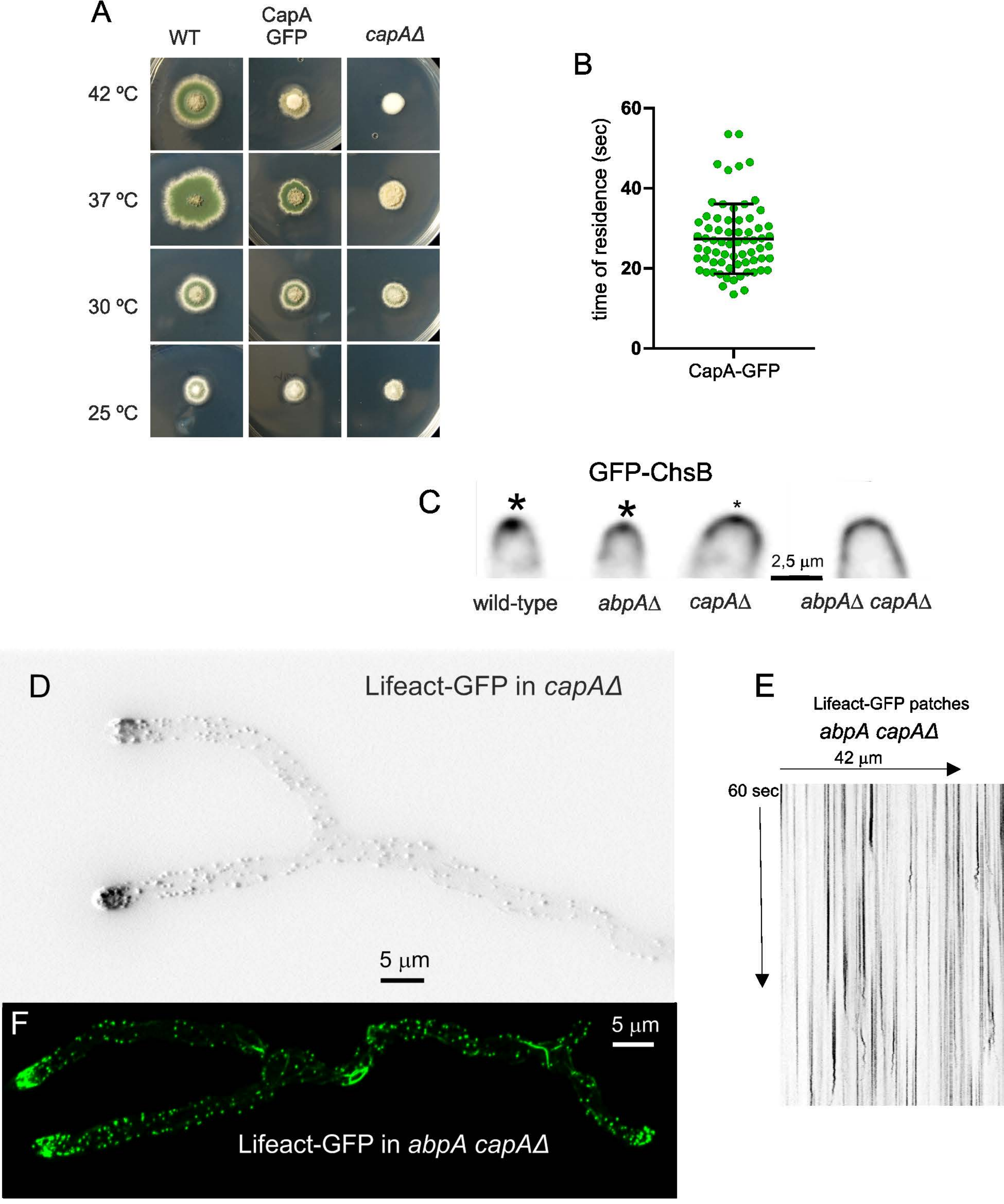

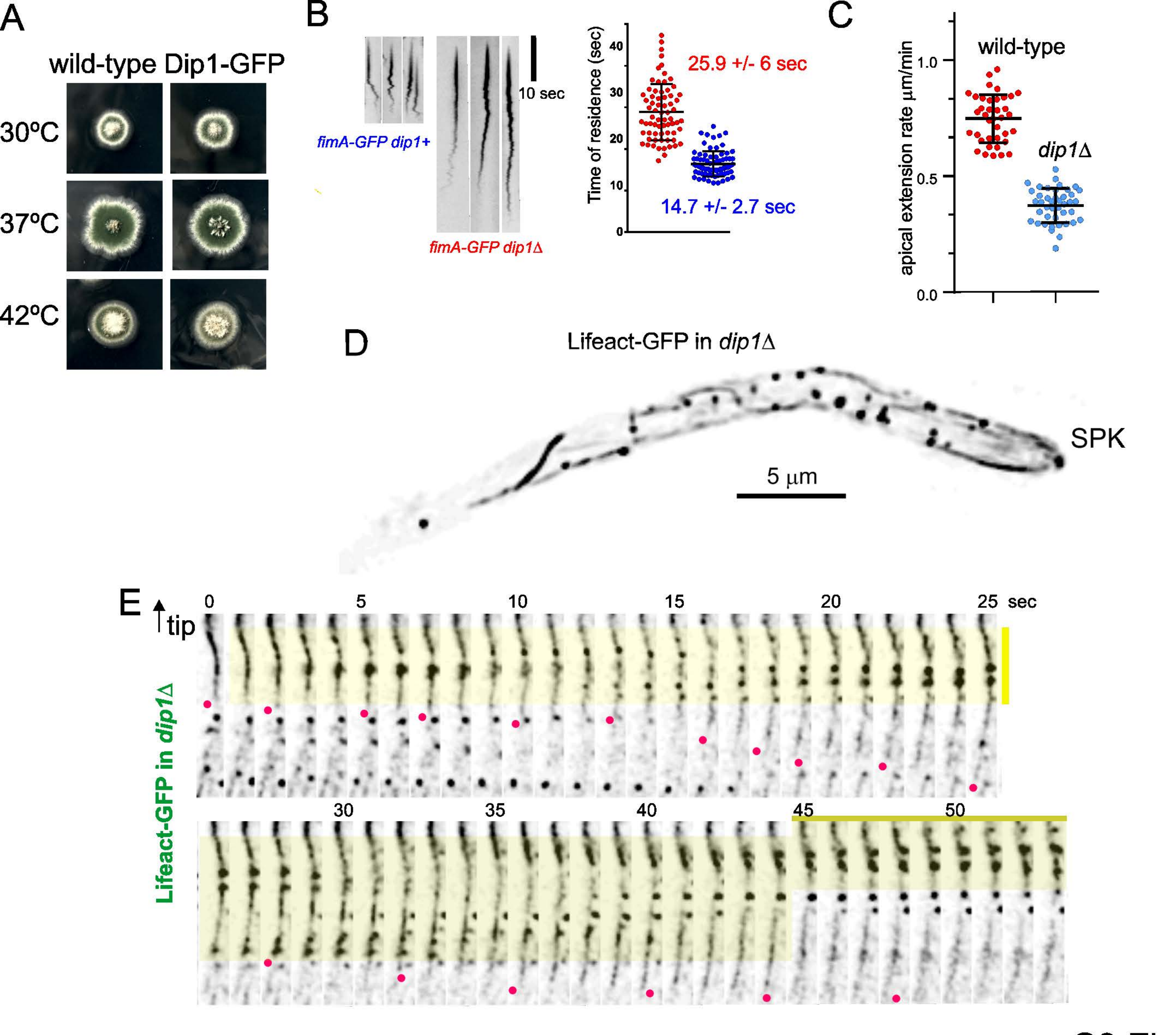

S10 Fig

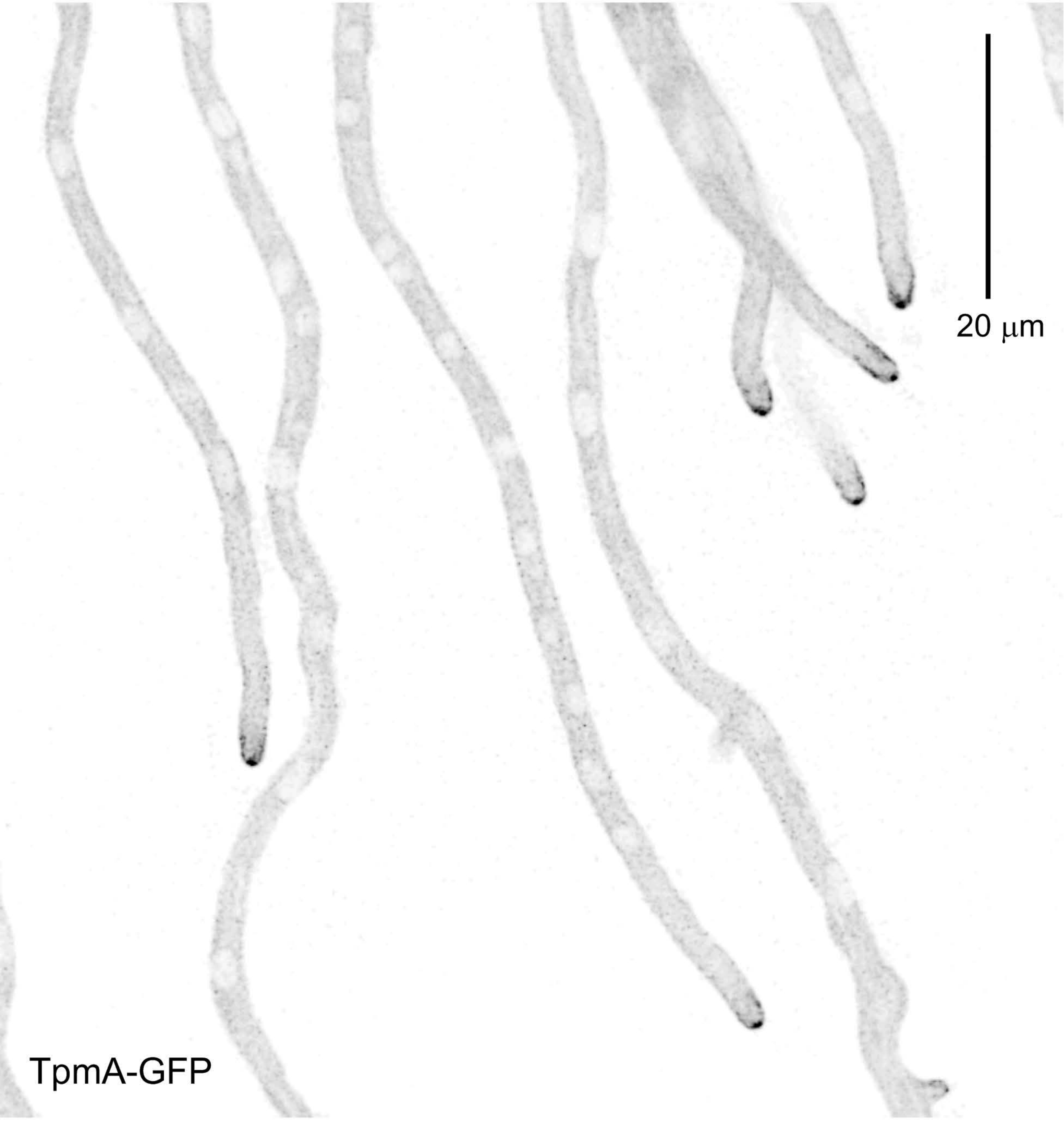
